# Rats use darting as a strategy to navigate between reward and safety during platform-mediated active avoidance under different social contexts

**DOI:** 10.64898/2026.05.21.726998

**Authors:** Karissa Payne, Shannon Ruble, Halle Ness, Helen Durrett, Cassandra Kramer, Maria M. Diehl

## Abstract

The platform-mediated active avoidance (PMA) task has been used as a rodent model of decision-based active avoidance in which rat learn to avoid a tone-signaled shock. Prior studies utilizing the PMA task have primarily investigated avoidance, freezing, and food-seeking behaviors, but few studies have thoroughly assessed darting behavior, a more recently identified measure of fear that has been largely explored in conditional fear paradigms. Here, we investigated the properties of darting that occur during the PMA task, in which rats either acquired the PMA task alone or with a social partner. We found that rats undergoing solitary PMA produced significantly more darting bouts, whereas rats undergoing social partner PMA produced darts that were faster and shorter in duration. We also found that darting in solitary PMA was predominantly concentrated at the platform, whereas darting in social partner PMA occurred more often outside of the platform and lever zones. Analysis of darting trajectories, which included movements surrounding each darting bout, revealed that darting was embedded in a broader movement strategy between the platform and lever zones, especially during solitary PMA, and this pattern increased across training days. These findings suggest that darting during the PMA task serves as a learned strategy to navigate between reward and safety and is modulated by social context, which is distinct from escape-like darting observed in auditory fear conditioning.

## Introduction

The platform mediated avoidance (PMA) task was first developed as a modification of auditory fear conditioning, in which the subject learns to associate a tone stimulus with an aversive footshock unconditioned stimulus (US) and must endure the shock US. The PMA task differs by allowing subjects to adopt a strategy to avoid the impending shock US by stepping onto a safe platform (Bravo-Rivera et al., 2014). The PMA task enabled researchers to study an active response to danger, in contrast to the reactive response of freezing that had been commonly used as a measure of fear learning for decades (Diehl et al., 2019, 2024; Sangha et al., 2019). Moreover, the PMA task allows subjects to lever press for a sucrose reward, creating a motivational conflict between reward seeking and threat avoidance. The PMA task, therefore, has become an ideal task to study realistic situations of active avoidance, in which individuals often must forego an appetitive stimulus to protect themselves from potential threats (Diehl et al., 2019).

Much of the research using the PMA task has assessed freezing during the tone and suppression of lever pressing for reward to compare aversive learning between the PMA task and auditory fear conditioning. However, recent studies on fear conditioning have revealed a behavioral strategy called darting, an escape-like response that emerges during conditioning (Colom-Lapetina et al., 2019; Gruene et al., 2015). Many of these studies found that darting responses were more common in female rats, compared to male rats (Colom-Lapetina et al., 2019; Gruene et al., 2015; Manzano Nieves et al., 2023; Mitchell et al., 2024; Shansky, 2004). Subsequent work demonstrated that external factors such as the size of the operant chamber, shock intensity and other parameters could reveal darting in both male and female rats (Mitchell et al., 2022).

Darting behavior has since been reported across multiple aversive learning paradigms including social threat conditioning (Lozano-Ortiz et al., 2025), discrimination of fear, safety, and reward (Greiner et al., 2019), and active avoidance (López-Moraga et al., 2025; Ruble et al., 2025). In contrast to darting during fear conditioning, darting during these tasks is not sex-specific. This discrepancy in findings suggests that darting may be a complex adaptive strategy that serves different purposes in fear conditioning versus other behavioral paradigms. We were therefore interested in investigating the spatial and temporal dynamics of darting in PMA to further understand how and when this behavior arises.

The present study therefore examined the temporal, kinematic, and spatial properties of darting behavior during solitary and social partner PMA training. We quantified dart frequency, speed, duration, distance, and spatial trajectory organization across training and examined how darting related to freezing, platform avoidance, and lever pressing behaviors. We hypothesized that darting would increase across days because previous studies have shown that darting is a learned response (Greiner et al., 2019; Gruene et al., 2015; Le et al., 2024) much like active avoidance. We also hypothesized that darting speed, distance, or duration might differ by social condition or be sex-dependent since darting frequency alone during the PMA task had not previously shown any sex differences (Ruble et al., 2025). We also investigated the location of darting between the food-seeking and safety-seeking areas of the task as PMA training progresses. We hypothesized that darting location would become more organized and concentrated around the platform as training progressed since these are the two most important areas in the operant chamber for the PMA task. This would also reflect darting as an active avoidance strategy, in which rats use darting as a navigational strategy to move between the reward and safety zones of the PMA task.

## Materials and Methods

### Subjects

104 Sprague Dawley rats (50 female, 54 male) were bred in-house from rats purchased from Charles River (Wilmington, MA) and same sex housed in groups of 2-3 rats per cage on a reverse light cycle and tested in the dark phase of their light cycle. Rats were handled and weighed twice weekly, or daily during experiments. When all rats reached the weight indicated by the standard growth chart for Sprague Dawley rats at approximately 8-10 weeks of age, they were placed on a restricted diet (16-18 g/day) of standard laboratory rat chow to facilitate lever pressing for sucrose pellets while still maintaining at least 85% of their target body weight. All procedures were approved by the Institutional Animal Care and Use Committee of Kansas State University in compliance with the National Institutes of Health guidelines for the care and use of laboratory animals.

### Behavioral Training

Rats were trained to press a lever to receive a sucrose pellet (BioServ, Flemington, NJ) across 5-7 days inside operant boxes, as previously described (Kramer et al., 2025; Ruble et al., 2025). Briefly, rats progressed through a fixed-ratio one (FR-1), followed by a 15 sec variable-interval (VI-15), and finally a VI-30 schedule of reinforcement. Once lever press training was completed, rats underwent PMA training and were conditioned with a pure tone (30 sec, 4 kHz, 75 dB) co-terminating with a footshock (2 sec, 0.4 mA). Rats received 9 tone-shock pairings per day for 10 days, with an inter-trial interval (ITI) averaging 3 min while maintaining a VI-30 schedule throughout PMA training. The availability of reward on the opposite side of the platform motivated rats to leave the platform during the ITI, facilitating trial-by-trial assessment of avoidance.

For solitary PMA, rats underwent training alone in standard conditioning chambers (Coulbourn Instruments). For social partner PMA, rats were assigned a same-sex and age-matched partner that was not a cage mate. Partner rats were placed one on either side of the perforated acrylic barrier, where they were still able to see, smell, and hear one another while undergoing 10 days of training simultaneously.

### Data Collection

Behavioral data used in the present analyses were previously collected and partially reported in (Ruble et al., 2025) and were subsequently curated and re-analyzed using updated behavioral classification and statistical procedures. ANY-Maze software was used to detect the animal’s location, movements, and behaviors throughout PMA training. Several zones within the operant chamber were designated to map the rat’s movement. A rat was considered within a zone when their center of body mass crossed into the zone. Freezing behavior was detected using ANY-maze. A minimum freeze duration of 250 ms was required to be calculated as a freezing bout. Darting bouts were classified as a brief (300-1000 ms) and quick movement (at least 23.5 cm/s) occurring during the tone period while the rat was not receiving shock. Darting bouts were considered to be separate events if they were separated by 0.8 s or more, to avoid double-counting a single locomotor bout. Darting location was quantified using x-y coordinates of the video recording. All behavioral measures were collected across the 10 days of training in solitary and social partner PMA in male and female rats.

Four quality control filters were applied to remove non-darting movements. Segments (n=4566) exceeding 1 s in duration were excluded, as these likely reflected running or exploratory movements rather than darting. Segments (n=7) with a maximum speed exceeding 3.6 m/s were excluded, as these surpass speeds physiologically plausible for rats locomoting voluntarily within a confined operant chamber, and are more likely to be tracking artifacts than actual movements. Finally, segments (n=1987) that overlapped with shock delivery were excluded unless the animal was on the avoidance platform during the shock. Movements accompanying an experienced shock were classified as reflexive escape responses rather than proactive anticipatory darts. Finally, any darting bouts where AnyMaze did not record a difference in x-y coordinates between the beginning and end of the dart were not counted, as those are also presumed to be a tracking error. This left a total of n=46,502 segments that were analyzed.

Following dart identification, a ±1 second trajectory window was extracted around each darting bout (1 s before the onset and 1 s after the offset), capturing the approach (PreDart), the dart itself (MidDart), and the post-dart trajectory (PostDart). Behavioral measures quantified across training included darting frequency, dart speed, dart duration, dart distance, freezing behavior, time spent on the avoidance platform, lever pressing behavior, and spatial distribution of dart trajectories across ROIs.

### Data Analysis

Behavioral data were analyzed using multilevel generalized linear mixed-effects models (GLMMs) to account for repeated measurements across training days and tone presentations, non-normal outcome distributions, and subject-level variability. Multilevel modeling approaches are well suited for repeated-measures behavioral datasets with nested observations, unequal variance structures, and missing or excluded data points (Bolker, 2015; Sommet & Morselli, 2017). Multilevel regressions were performed on each behavioral measure of interest to assess differences in darting behavior observed during PMA training. For dart speed and dart distance, Gamma generalized linear mixed models (GLMMs) with a log link were performed using the glmmTMB package (Brooks et al., 2017) to account for the continuous, positively skewed, and strictly positive nature of these measures. For dart duration, the same Gamma GLMM approach was applied. For all three continuous dart metrics, Condition (social/solitary), Sex, and Training Day were included as fixed effects (with all two- and three-way interactions), and random intercepts were fitted for individual subjects. For number of darting bouts per tone, a multilevel negative binomial GLMM (similar to Poisson regression for count data, but accounting for overdispersion) (Gardner et al., 1995) was performed using the glmer.nb function in the lme4 package (Bates et al., 2014), with the same fixed and random effect structures as were used in the models for dart speed and distance.

To investigate whether measures of freezing, avoidance, or lever pressing predicted darting behavior, three additional negative binomial GLMMs were fitted using lme4, with darting bout count as the outcome and percent time freezing, percent time on the platform, or number of lever presses as continuous predictors, alongside condition (social/solitary), sex, and training day (scaled 0–1 to help with model convergence) and all interactions.

To characterize the spatial trajectory of darting bouts relative to apparatus zones (lever zone, platform zone, both, or neither, with each zone being compared against the neither or “non-zone” darting category), a Bayesian multinomial GLMM was fitted using the brms package (Bürkner, 2017), with zone-entry outcome (none, lever, platform, or both) as the categorical response variable and Condition, Sex, Training Day, and Tone Number (with all interactions) as fixed effects, and individual subject random intercepts. For these Region of Interest (ROI) analyses, we used a multinomial model to estimate the likelihood of entering all ROIs simultaneously, and we used a multilevel structure because the data consisted of repeated observations nested within participants. A Bayesian estimation framework was used for these models, because the combination of multilevel and multinomial components creates a statistically complex model that is difficult to estimate using traditional methods. Bayesian methods provide a flexible approach for estimating these parameters while appropriately accounting for uncertainty in the model (Kass & Raftery, 1995; Kruschke, 2015). Bayesian inference quantifies the probability of the observed data, given a range of possible parameter values, unlike traditional means (i.e. frequentist statistics), which evaluate evidence against a null hypothesis and express uncertainty as a p-value. Results from these models are expressed as posterior means with 95% credible intervals (CI), which is the range in which the parameter value falls within 95% probability and as directional Bayes Factors (BF) (Makowski et al., 2019), which reflect the relative evidence in favor of an effect, compared to a null hypothesis of no difference.. A BF of 20 or greater indicates strong evidence for the tested effect (Jarosz & Wiley, 2014). Model random-effects structures were compared using WAIC values, and the best-fitting model was selected for inference. Post-hoc contrasts and estimated marginal means were computed using the emmeans package (Lenth et al., 2022) with Tukey adjustment where applicable. All analyses were conducted in R (version 4.4.2; R Core Team, 2026), using glmmTMB (Brooks et al., 2017), lme4 (Bates et al., 2014), brms (Bürkner, 2017), and emmeans (Lenth et al., 2022).

## Results

### Darting frequency is greater during solitary PMA, whereas darting speed is greater during social partner PMA

Our prior work reported that, similar to auditory fear conditioning, rats exhibited darting during the PMA task (Ruble et al., 2025), and we were interested in how certain features of darting might differ between solitary and social partner PMA. To characterize darting behavior during PMA, we measured frequency, speed, duration, and distance of the darting behavior across the 10 days of training during the solitary and social versions of the PMA task in male and female rats. **Figure 1A** shows example velocity traces of a rat during the solitary PMA task (blue) and the social partner PMA task (purple), along with examples of dart and non-dart movements at early and late stages of solitary PMA (**Figure 1A**, right).

**Figure 1:**
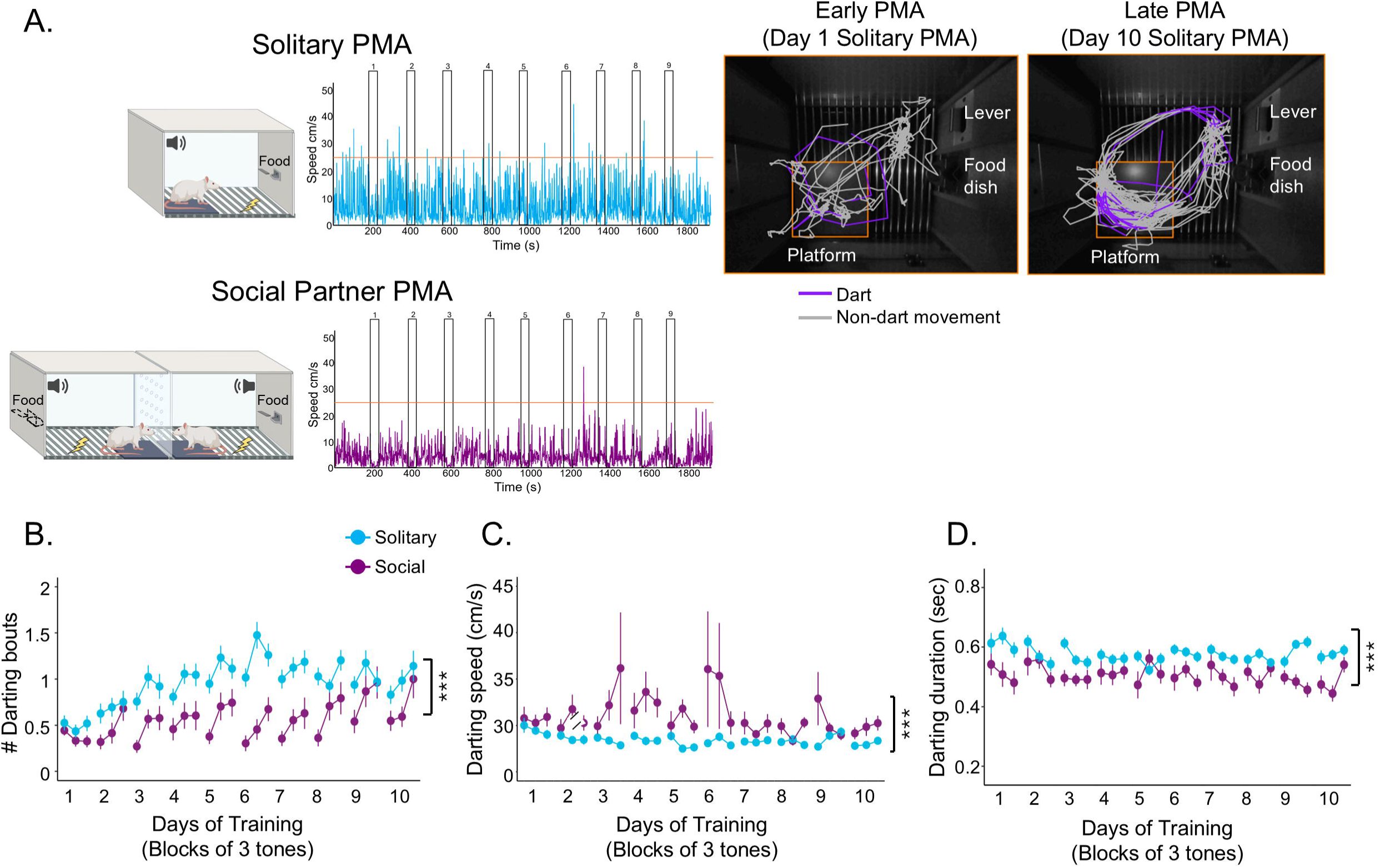
Darting properties and kinematics. **(A)** *Left*: Schematic of PMA task and velocity traces from individual rats during a single session of the Solitary (*top,* blue, n=59) and Social Partner (*bottom,* purple, n=45) PMA tasks. Orange line in velocity traces indicate the threshold speed for a dart (23.5 cm/s) and tones are indicated by white bars. Right: Trajectories of an individual rat showing the location of dart (purple) vs. non-dart (grey) movements during the 9 tones during early vs. late days of solitary PMA training. **(B)** Number of darting bouts, **(C)** darting speed, **(D)** and darting duration across 10 days of PMA training in social partner (purple) or solitary PMA (blue). Solitary PMA rats performed more darting bouts (z=-4.400, *p*<0.001) and had longer darting durations (z=-4.084, *p*<0.001) than social partner PMA rats. Social partner PMA rats exhibited increased darting speed (z=3.851, *p*<0.001) compared to solitary PMA rats. Data shown as mean ±SEM. ***p<0.001.

A multilevel negative binomial regression revealed that rats in the solitary PMA task produced significantly more darting bouts than rats in the social partner PMA task across training (z = -4.40, *p* < 0.001; **Figure 1B**; see **Supplementary Table 1** for parameter estimates). In addition, the number of darting bouts significantly increased across training days (z = 4.539, *p* < 0.001). No significant main effect of sex was observed (z = 0.236, *p* = 0.813), and no significant two- or three-way interactions were detected (all *p*’s > 0.05; **Supplementary Figure 1A**). When separated by sex, both females (z = −4.843, *p* < 0.001) and males (z = −3.570, *p* < 0.001) exhibited significantly fewer darting bouts during social partner compared to solitary PMA, confirming that condition differences are not driven by a single sex.

To characterize the kinematic properties of darting, we next assessed darting speed, distance, and duration across all training days. A multilevel gamma regression revealed that darting speed in social partner PMA was significantly greater than in solitary PMA (z = 5.541, *p* < 0.001; **Figure 1C**; see **Supplementary Table 2** for parameter estimates). In addition, darting speed significantly decreased across training days (z = −2.643, *p* = 0.008). There was no significant main effect of sex (z = 0.479, *p* = 0.632) or significant two- or three-way interactions (all *p’s* > 0.05; **Supplementary Figure 1B**). When separated by sex, both females (z = 6.220, *p* < 0.001) and males (z = 6.790, *p* < 0.001) exhibited significantly greater darting speed during social partner compared to solitary PMA again confirming that condition differences were not sex-dependent.

A multilevel gamma regression revealed that rats in social partner PMA exhibited significantly shorter darting durations compared to rats in solitary PMA (z = −4.084, *p* < 0.001; **Figure 1D**; see **Supplementary Table 3** for parameter estimates). No significant main effects of sex (z = 0.787, *p* = 0.432) or Day Number (z = −0.609, *p* = 0.543) were observed, and no significant two- or three-way interactions were detected (all *p’s* > 0.05; **Supplementary Figure 1C**). When separated by sex, both females (z = −7.080, *p* < 0.001) and males (z = −5.164, *p* < 0.001) exhibited significantly shorter dart durations during social compared to solitary darting, again confirming that neither sex was independently driving the condition effect.

A multilevel gamma regression examined darting distance showed that darting distance significantly decreased over days (z = −2.930, *p* = 0.003; see **Supplementary Table 4** for parameter estimates). No significant main effects of Condition (z = 1.535, *p* = 0.125) or sex (z = −0.102, *p* = 0.919) were observed, and no significant two- or three-way interactions were detected (all *p’s* > 0.05). Comparisons between females and males within each condition revealed no significant sex differences in social partner PMA darting distance (z = −1.288, *p* = 0.198) or solitary PMA darting distance (z = 1.080, *p* = 0.280). Taken together, these results show that for active avoidance, darting is more frequent under solitary conditions, and the speed at which rats dart is faster and the duration shorter under social conditions. In contrast to fear conditioning studies reporting sex differences in darting frequency, we did not find that darting behavior or any properties of darting were sex-dependent.

### Darting occurred most often at the platform and became more concentrated at the platform as PMA training progressed

During fear conditioning, subjects learn the tone shock association and exhibit a physiological response to the tone CS, whereas during the PMA task, subjects also learn an adaptive strategy to avoid the shock US by moving to the safe platform while foregoing the opportunity to obtain a food reward (Diehl et al., 2018, 2019, 2024; Sangha et al., 2019). Because the context and motivation in the PMA task greatly differs from auditory fear conditioning, we were interested in how rats may be using darting behavior as a strategy to navigate between the reward and safety zones (lever and platform areas, respectively) during the PMA task. We observed that rats change their dart and non-dart movements as training progresses in the PMA task (**Figure 1A**, right). Therefore, we wanted to determine if the location of darting differed by task type (solitary vs. social partner), training day, or sex. To do this, we assessed the location of darting within the operant chamber. Because of the complex nature of modeling multiple potential ROIs across multiple subjects, we used a Bayesian multinomial regression to classify each darting bout by its spatial location (see Data analysis section in the Methods). **Figure 2A-B** shows heat maps of darting locations in female (left columns) and male (right columns) at early PMA training (top row) and late PMA training for solitary (**Figure 2A**) and social partner (**Figure 2B**) PMA tasks. Dart locations were defined as occurring in the platform zone, lever zone, both zones (darting between them), or neither zone (**Figure 2C**).

**Figure 2:**
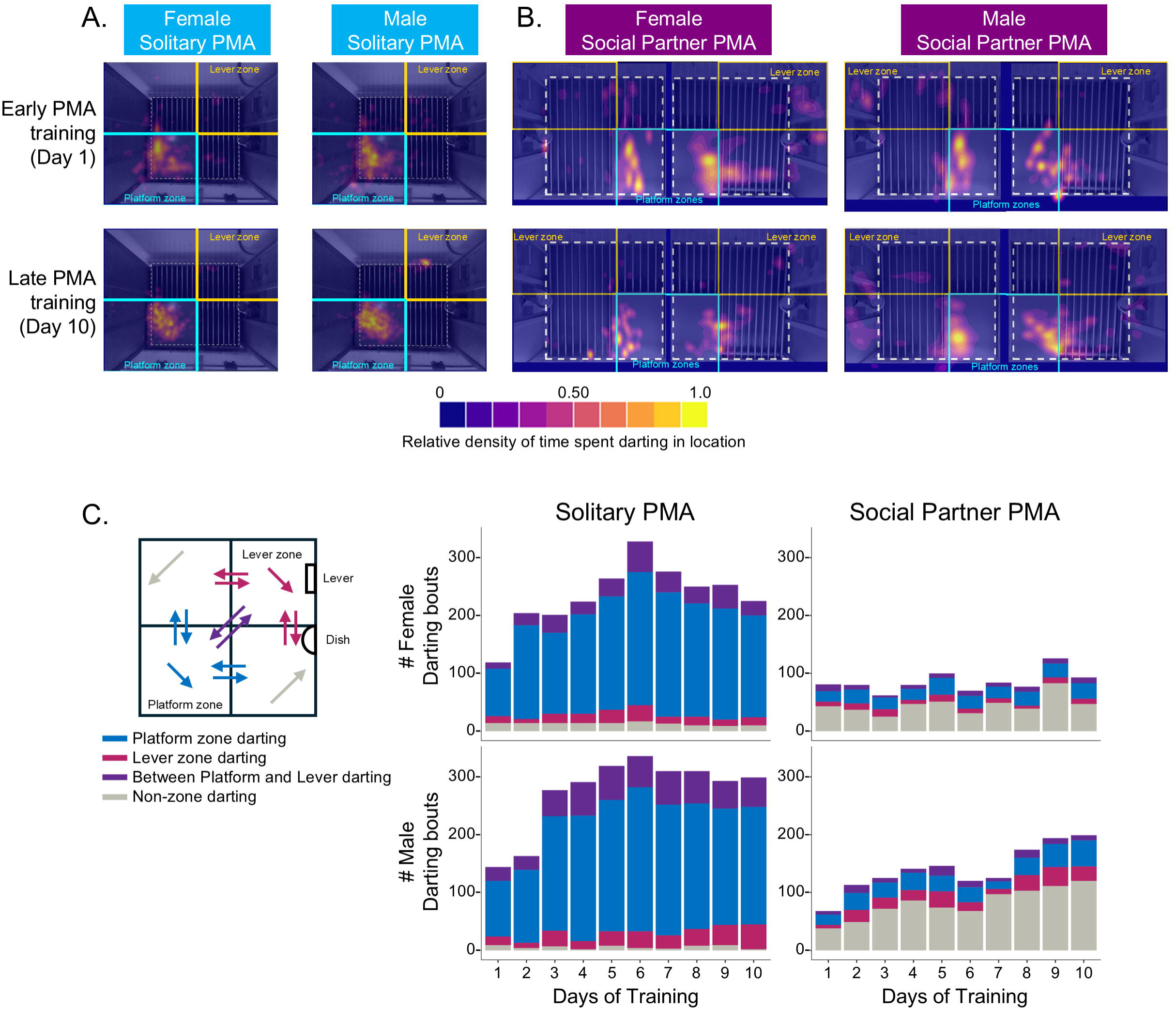
Darting occurs more frequently in the platform or between platform and lever zones in solitary PMA, whereas most darting occurs outside either zone in Social Partner PMA. **(A)** Heat maps showing location of darting during solitary PMA on day 1 (*top*) and day 10 (*bottom*) during the tone, also separated by females (*left*) and males (*right*). **(B)** Heat maps showing location of darting bouts during social partner PMA on day 1 (*top*) and day 10 (*bottom*) during the tone, also separated by females (*left*) and males (*right*). **(C)** Quantification of darting location: darting in the platform zone (blue), lever zone (pink), darting between platform and lever zones (purple), and darting outside of either zone (grey) during the tone period. Darts were substantially more likely to occur in the platform zone compared to other zones during solitary PMA (*middle*, 95% CI [0.678,0.878]) whereas darts were more likely to occur in the non-zone during social partner PMA (*right*, 95% CI [0.458, 0.673]). There were no sex differences in spatial dynamics of darting in solitary PMA (middle). In social partner PMA, the probability of non-zone darting increased (*gray*, 95% CI [0.007, 0.046]) and the probability of platform zone darting decreased (*blue,* 95% CI [-0.035, -0.003]) in male rats (*bottom right*) whereas the probability of lever-zone darting decreased (*pink,* 95% CI [-0.020, -0.002]) in female rats (*top right*).

A Bayesian multilevel multinomial regression showed strong differences in the spatial distribution of darting behavior between social partner and solitary PMA, with comparatively little evidence for sex effects. Compared to darts that occurred in neither zone (“non-zone” darting), darts in social partner PMA were substantially less likely to occur in the platform zone (B = - 1.670, 95% CI [-2.189, -1.193], BF > 1000), the lever zone(B = -0.743, 95% CI [-1.307, -0.213], BF = 332.33), or between the platform and lever zones (B = -1.237, 95% CI [-1.721, -0.772], BF > 1000; **Figure 2C**; see **Supplementary Table 5** for parameter estimates and comparisons). Social partner darting also showed negative interactions with Training Day across all zones, indicating decreasing zone-associated darting over training days, particularly for the platform zone and between platform and lever zones. Sex effects and higher-order interactions were generally weak, with credible intervals (CIs) overlapping zero and BFs mostly favoring weak or anecdotal evidence.

Posterior estimated probabilities indicated that social partner darting was most likely to occur outside the platform and lever zones (“non-zone”; M = 0.568, 95% CI [0.458, 0.673]), whereas solitary darting overwhelmingly occurred in the platform zone (M = 0.791, 95% CI [0.678, 0.878]). Direct contrasts confirmed that social partner darting was substantially more likely than solitary darting to occur in neither zone (difference = 0.546, 95% CI [0.435, 0.651], BF > 1000), while solitary darting was substantially more likely to occur in the platform zone (difference = -0.530, 95% CI [-0.678, -0.360], BF > 1000). There was little evidence for differences between solitary and social partner PMA in the “both” category (between platform and lever zones) or the lever zone.

Sex-specific posterior means showed highly similar spatial profiles for females and males. For both sexes, darting in the social partner PMA task was characterized by elevated probabilities of occurring outside either the platform or the lever zones, whereas darting in the solitary PMA task remained dominated by platform-associated bouts. Within females, darting during social partner PMA was more likely to occur outside the lever or platform zones compared to solitary PMA (difference = 0.515, 95% CI [0.357, 0.659], BF > 1000) and less likely to occur in the platform zone (difference = -0.529, 95% CI [-0.726, -0.283], BF > 1000). The same pattern was present in males, with nearly identical effect sizes. No credible sex differences emerged within either darting type. This supports that the condition differences were not driven by either sex.

Training Day slope estimates suggested modest temporal changes in darting location in social partner but not solitary PMA. In social partner PMA, males showed increasing probability of “non-zone” darting across days (Estimate = 0.0263, 95% CI [0.0071, 0.0455], BF = 269) and decreasing probability of platform-associated darting (Estimate = -0.0173, 95% CI [-0.0353, - 0.0025], BF = 97). Females in social partner PMA showed decreasing lever-associated darting across training days (Estimate = -0.0089, 95% CI [-0.0204, -0.0015], BF = 114). In solitary darting, day-related effects were comparatively small. Overall, these findings indicate that social partner PMA darting increasingly shifted away from zone-defined spatial locations over training, whereas solitary PMA darting remained strongly concentrated within the platform zone.

The above analysis revealed that the dart itself largely occurs in and around the platform, and not predominantly between the platform and lever zones. Because we observed rats moving exclusively between the platform and lever during the tone, especially in late stages of PMA (see **Figure 1A**), we therefore reasoned that there might also be some movement occurring immediately before and after the dart that would indicate rats are using darting behavior together with non-darting movement to navigate between the food reward and platform. We termed this a “darting trajectory.” Another Bayesian multinomial regression was performed that included behavioral data from 1 sec preceding and following each darting bout, to examine darts as part of a larger movement trajectory. **Supplementary Figure 3A** shows examples of this larger movement darting trajectory, with 1 sec of movement before the dart (PreDart, light pink), the dart itself (MidDart, purple), and 1 sec of movement after the dart (PostDart, dark purple) across Solitary (top) and Social Partner (bottom) PMA training.

For darting trajectories occurring in the lever zone relative to non-zone regions, there was strong evidence for an effect of Training Day when lever zone darting was compared to non-zone darting (B = 0.523, 95% CI [0.175, 0.958], BF > 1000; **Supplementary Figure 3B**), indicating that lever-associated darting increased across training. There was also strong evidence for a Condition × Day Number interaction (B = −0.451, 95% CI [−0.879, −0.126], BF = 726), suggesting that this increase in lever zone darting across training was reduced in social partner compared to solitary PMA. No substantial evidence was observed for main effects of Condition or Sex, or for higher-order interactions involving Sex (all BFs < 10). There was also strong evidence that darting trajectories in the platform zone was less likely in social partner compared to solitary PMA (B = −2.114, 95% CI [−3.396, −0.925], BF > 1000). Strong evidence also indicated that platform-associated darting increased across training days (B = 0.491, 95% CI [0.174, 0.914], BF > 1000), alongside a strong negative Condition × Training Day interaction (B = −0.515, 95% CI [−0.933, −0.200], BF > 1000), indicating that the increase across days was attenuated during social partner PMA.

There was decisive evidence that the likelihood of darting trajectories occurring in both zones compared to the non-darting zone (B = −2.248, 95% CI [−3.514, −1.063], BF > 1000) decreased during social partner PMA. Strong evidence also demonstrated increases in darting trajectories between the lever and platform zones across training days (B = 0.502, 95% CI [0.189, 0.920], BF > 1000), as well as a negative Condition × Day Number interaction (B = −0.508, 95% CI [−0.925, −0.194], BF > 1000), such that darts that traversed both the lever area and the platform declined across training in social but not solitary rats. Posterior estimated marginal means indicated that darting trajectories were substantially more likely to occur outside of both the lever and platform zones in social partner compared to solitary PMA. In contrast, darting trajectories were more likely to occur in both zones during solitary PMA. Posterior contrasts demonstrated near-complete posterior certainty that darting trajectories were more likely to occur in neither zone (“non-zone”; BF > 1000) or in the lever zone (BF > 1000) during social partner compared to solitary PMA. Darting trajectories were less likely to occur in both regions (BF = 9) during social partner compared to solitary PMA. No credible difference between social partner and solitary darting was observed for platform-only darting trajectories.

When separated by sex, darting trajectories were more likely to occur in neither zone or in the lever zone in social partner compared to solitary PMA in both females and males. Males additionally showed strong evidence that darting trajectories were less likely to occur in both zones in social partner compared to solitary PMA (BF = 28). The corresponding effect in females was weaker and less certain (BF = 6). No credible sex differences in zone-specific darting trajectory probabilities were observed within either social partner or solitary PMA, as all posterior contrasts included zero within their credible intervals. Finally, posterior Training Day trend analyses revealed little evidence for meaningful changes across training in region-specific darting probabilities within sex or condition subgroups, as nearly all credible intervals overlapped zero. Taken together, these findings reveal that darting during PMA is spatially structured around the safety zone of the platform, especially in solitary PMA, and that this spatial structure emerges and becomes refined as training progresses. Whereas solitary rats used darting more for safety seeking, social partner rats used darting more outside of the safety and reward areas of the task, suggesting that social context may alter the spatial structure underlying darting behavior.

### Darting behavior was negatively correlated with freezing and avoidance and positively correlated with lever pressing

In our prior study, we found that females showed more avoidance compared to males in solitary PMA, and across both solitary and social partner PMA, males lever pressed more than females (Ruble et al., 2025). We also found no sex differences in freezing in either version of the task. To examine the functional relationship between darting and the above behavioral measures acquired during PMA, we determined whether freezing, time on platform, and lever pressing were reliable predictors of darting during the tone period. Greater freezing was associated with significantly fewer darting bouts overall (z = −13.939, *p* < 0.001; **Figure 3A**; see **Supplementary Table 7** for parameter estimates). No significant main effects of sex (z = −0.326, *p* = 0.745) or Training Day (z = 0.011, *p* = 0.991) were observed. A significant Freeze × Sex × Training Day interaction indicated that the negative relationship between freezing and darting strengthened across training days (z = −2.155, *p* = 0.031). A significant four-way interaction further qualified this effect: this day-dependent freeze-dart tradeoff was most pronounced in social partner PMA females (z = −2.672, *p* = 0.008), suggesting that among female rats in Social PMA, freezing and darting became increasingly mutually exclusive behaviors as training progressed. No other two- or three-way interactions reached significance (all *p* > 0.05). Slope analyses demonstrated that greater freezing predicted fewer darting bouts in both social partner (slope = −3.269, *p* < 0.001) and solitary PMA (slope = −3.571, *p* < 0.001), with no significant difference in freezing-related slopes between conditions (z = 0.764, *p* = 0.445). However, the negative relationship between freezing and darting was significantly stronger in females than males (z = −2.672, *p* = 0.0075).

**Figure 3:**
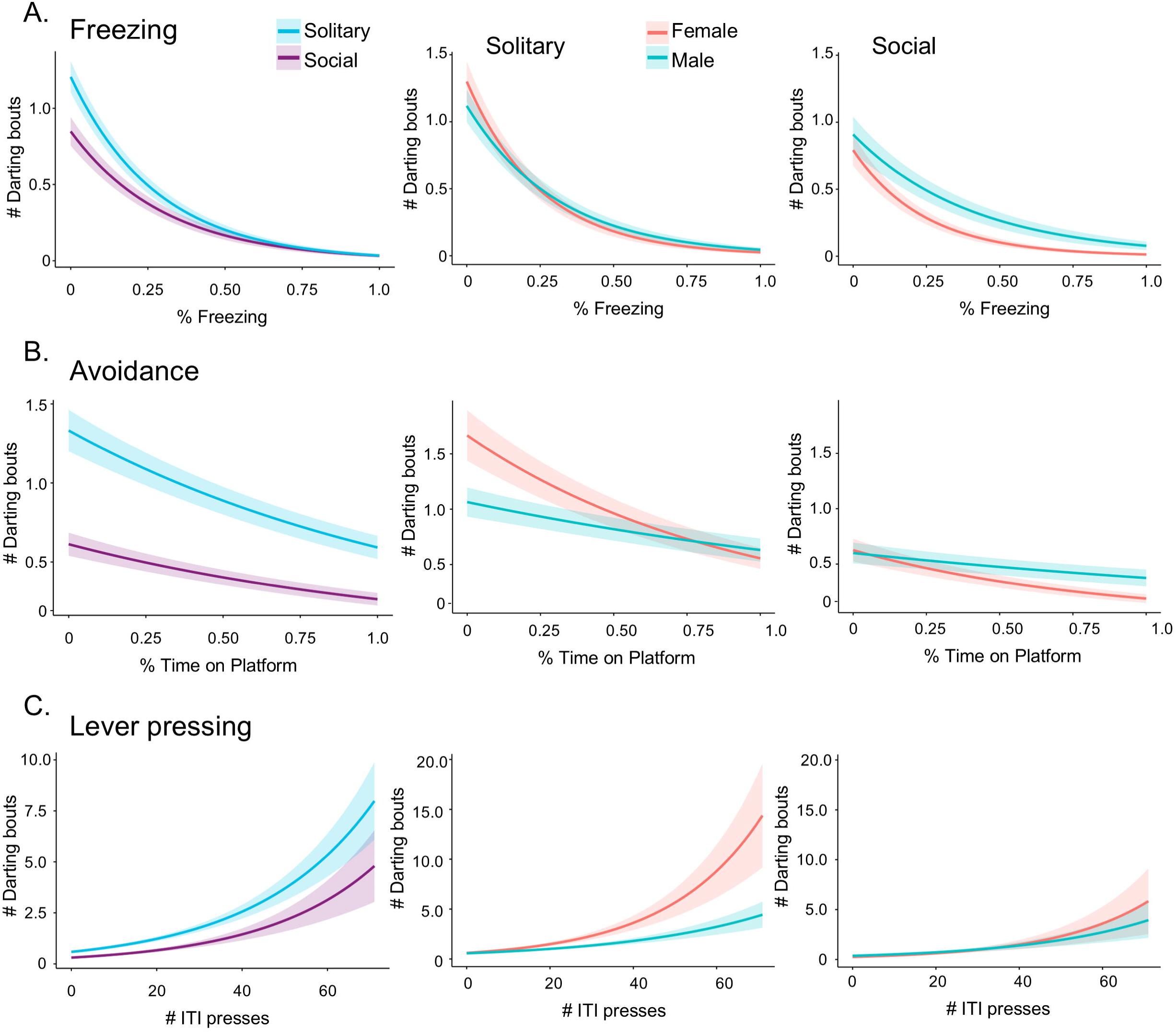
Darting is negatively correlated with freezing and avoidance and positively correlated with lever pressing for food reward. **(A)** Conditional effects plots showing whether tone-induced freezing predicts darting frequency when comparing rats in solitary (*blue*) vs. social partner (*purple*) PMA (*left*), female (*salmon*) and male (*teal*) rats in solitary PMA (*middle*), and female and male rats in social partner PMA (*right*). Increased freezing was associated with decreased darting (z= -13.939, *p*<0.001). The negative relationship between freezing and darting was significantly stronger in females compared to males (z= -2.981, *p*=0.003) regardless of condition. **(B)** Conditional effects plots showing whether time on platform during the tone predicts darting frequency in the same comparisons as **(A)**. Increased avoidance was associated with decreased darting (z=-7.142, *p*<0.001). The negative relationship between avoidance and darting was significantly stronger in females compared to males (z=-3.912, *p*<0.001) regardless of condition. **(C)** Conditional effects plots showing whether the number of lever presses during the ITI predicts darting frequency in the same comparisons as **(A)** and **(B)**. Increased lever pressing was associated with increased darting (z=11.369, *p*<0.001). The positive relationship between lever pressing and darting weakened across training (z=-5.707, *p*<0.001). There were no significant sex differences in the strength of the positive relationship between lever pressing and darting (z=1.859, *p*=0.063). Data shown as model fit line with error ribbons representing SE.

Greater avoidance, measured as the percentage of time spent on the platform, was associated with significantly fewer darting bouts overall (z = −7.142, *p* < 0.001; **Figure 3B**; see **Supplementary Table 8** for parameter estimates). A significant Avoid Percent × Sex × Day Number interaction emerged (z = −2.955, *p* = 0.003), whereas no significant four-way interaction between Avoid Percent, Condition, Sex, and Day Number was detected (z = −1.103, *p* = 0.270). No other two- or three-way interactions reached significance (all *p’s* > 0.05). Slope analyses demonstrated that greater avoidance predicted fewer darting bouts in both social partner (slope = −0.840, p < 0.001) and solitary PMA (slope = −0.810, *p* < 0.001), with no difference in avoidance-related slopes between conditions (z = −0.173, *p* = 0.863). However, the negative relationship between avoidance and darting was significantly stronger in females than males (z = −3.912, *p* < 0.001). Taken together, these results indicate that greater avoidance predicts fewer darting bouts across both versions of the PMA task, but the strength of this trade-off is significantly greater in females than in males.

Greater lever pressing was associated with significantly more darting bouts overall (z = 11.369*, p* < 0.001; **Figure 3C**; see **Supplementary Table 9** for parameter estimates). No significant main effect of sex was observed (z = 1.053, *p* = 0.293). Significant interactions emerged between Lever Count and Training Day (z = −5.707, *p* < 0.001), indicating that the relationship between lever pressing and darting weakened across training, as well as between sex and Training Day (z = −2.980, *p* = 0.003). No significant three- or four-way interactions involving Lever Count, Condition, Sex, and Training Day were detected (all *p* > 0.05). Slope analyses demonstrated that greater lever pressing predicted more darting bouts in both social partner (slope = 0.0387, *p* < 0.001) and solitary PMA (slope = 0.0367, p < 0.001), with no significant difference in lever-press-related slopes between conditions (z = 0.279, *p* = 0.781). Lever pressing also positively predicted darting in both females (slope = 0.0442, *p* < 0.001) and males (slope = 0.0312, *p* < 0.001), with no significant sex difference in slopes (z = 1.859*, p* = 0.063). In summary, lever pressing and darting co-occur early in training, suggesting that rats use darting to access reward quickly when they are less certain of when the shock may occur. As training progresses, however, this relationship begins to weaken as rats master the task and develop a more efficient strategy of balancing their motivational drives for safety and reward.

## Discussion

The current study found that more darting occurred in solitary compared to social partner PMA, darting was faster in social partner than in solitary PMA, and importantly, there were no sex differences in the quantity or location of darting bouts across PMA training. Most darting occurred at the platform, and to a greater degree during solitary than during social partner PMA. In fact, most of the darting bouts that were observed in social partner PMA occurred outside of the platform and lever zones, which was unexpected. When looking at movement before and after the dart bouts, termed darting trajectories, we found that rats were using darting as part of a larger movement to navigate between the lever and platform during the tone. Finally, we also found that darting bouts were negatively correlated with freezing and avoidance but positively correlated with lever pressing. There was a stronger negative relationship between both freezing and avoidance with dart frequency in females compared to males. Below, we discuss some caveats related to our findings and future directions.

The two major differences we noted between darting features in solitary and social partner PMA was that 1) darting bouts were faster in social compared to solitary PMA, and 2) dart location in social partner was outside of the safety and reward zones of the operant chamber. One possibility that may account for these findings may be related to the size of the operant chambers; the area of the social partner PMA chamber was smaller than the areas of the solitary PMA chamber. We reconfigured shuttle box chambers for our social partner PMA task (Kramer et al., 2025), which resulted in a slightly smaller area compared to the area available in the solitary PMA chambers. As a result, we also had to create a slightly smaller platform for the social partner PMA chambers so that rats could not remain on the platform while lever pressing for sucrose (Kramer et al., 2025). This configuration still allowed the same relative positions of the dish, lever, and platform while accommodating the smaller social chamber. In addition, the smaller area in which the rats could navigate between the lever and platform may have also led to their darting trajectories expanding out into the non-zone regions. Future studies may want to modify the social partner PMA task to include a larger space to determine if chamber size affects the speed and location of darting in the PMA task.

The findings reported here as well as previous PMA studies have not reported any sex differences in darting (Halcomb, et al., 2023; López-Moraga et al., 2025; Ruble et al., 2025), which notably contrasts with female-biased darting that has been reported in auditory fear conditioning (Colom-Lapetina et al., 2019; Gruene et al., 2015; Mitchell et al., 2024). This may be due to inherent differences in fear and avoidance tasks – the PMA task has added complexities involving instrumental responding, balancing the conflict between avoidance of shock and pursuance of reward. These may lead to the female bias for darting to disappear.

Despite not observing sex differences in the raw number of darts, we did find that females showed a stronger negative relationship between freezing and avoidance with darting frequency, suggesting that behavioral strategy choice is more likely to be exclusive in females whereas in males it is more flexible. When females freeze or avoid, they are less likely to switch to a darting strategy. On the other hand, males may show more flexibility between choosing behavioral strategies and seem more likely to switch between freezing, avoiding, and darting.

Furthermore, darting may be utilized in a different way for active avoidance tasks compared to passive fear tasks. During fear conditioning, darting is characterized as an escape-like reaction to the tone CS, sometimes replacing freezing as an expression of fear. During the PMA task, darting is spatially concentrated around the platform, suggesting that darting is being used as a navigational tool to obtain safety. As such, PMA-related darting may reflect a combination of fear-driven movement and instrumentally-reinforced avoidance of shock. Future PMA studies should consider assessing whether there are differences in escape response to determine if they might be correlated with darting bouts occurring exclusively during the tone CS.

The current study found interesting differences in darting related to the social context. Social partner PMA suppressed darting frequency while also increasing the speed and changing the location of darting during the task. This may reflect effects of social buffering if darting is occurring in a non-zone region that is outside of the lever and platform but closer to the social partner. Another possibility is that social partner rats may be darting in response to or in coordination with their partner’s movements. Future studies could classify one of the non-zone regions located near the social partner and determine if more darting or other behaviors are occurring in this zone compared to the same zone in solitary PMA. Other future directions could also assess social communication between the partners such as visual attention and ultrasonic vocalizations to determine their relationship to darting during PMA.

The nature of darting behavior as a learned versus non-associative behavior has been actively debated in the field, and the findings from the current study support the idea that darting is a learned behavioral strategy. One study reported that mice would display darting and other similar escape-like behaviors (jumping, running, etc.) to auditory stimuli or shock US simply because they were sudden changes in stimulation, not because darting became a learned response to a CS-predicting shock (Trott et al., 2022). The authors argued that darting reflects sensitization (similar to a startle response) rather than associative learning. Another study followed up on these findings and argued that although non-associative factors can contribute to the locomotor movement of darting, robust and sustained darting behavior requires explicit CS-US association (Le et al., 2024). The findings reported here agree with the latter idea because darting frequency during PMA increased systematically across training days and the location of darting also shifted to becoming more concentrated at the platform as training progressed. This interpretation is also consistent with prior work demonstrating that darting occurs primarily during the tone CS and increased across CS-US trials (Gruene et al., 2015), and that darting during a fear-reward-safety discrimination task is suppressed during the safety signal (Greiner et al., 2019).

The PMA task is an established rodent model of decision making under threat, with direct relevance to understanding the behavioral and neural mechanisms underlying neuropsychiatric disorders such as PTSD and other anxiety disorders (Diehl et al., 2018; Martínez-Rivera et al., 2023; Rodriguez-Romaguera et al., 2016). The darting behavior characterized here may offer an additional translational window into active coping strategies that are not captured by freezing or avoidance measures alone. In particular, the sex differences observed in the coupling between freezing and avoidance with darting bouts may be relevant to the higher prevalence of PTSD and other anxiety disorders in women, which has been linked to differences in active versus passive coping strategies (Cholankeril et al., 2023; Olff, 2017; Schneider et al., 2025). Future studies examining the neural substrates of darting during the PMA task, including circuits implicated in active avoidance such as prelimbic and infralimbic prefrontal cortex, basolateral amygdala, and ventral striatum (Bravo-Rivera et al., 2014, 2015; Diehl et al., 2019, 2020; Ruble et al., 2026) may reveal dissociable mechanisms underlying darting as a distinct active coping strategy.

## Author Contributions

Conceptualization, MMD; Research design, MMD; Methodology, MMD, KP; Investigation, SR, CK, HN, HD, MMD; Data analysis, KP, CK, HN, MMD; Supervision of experiments and analysis: SR, MMD; Writing, SR, KP, MMD; Funding Acquisition, MMD.

## Acknowledgements

We thank Drs. Susan Sangha and Abigail Polter for helpful comments on the manuscript. We also thank Tessa Maze, Jenna Thompson, Kelly Krehbiel, Ashvini Wickramasundara, Allison Moser, Allison Drouhard, Emma Wrampe, Charlotte Kettler, and Ivy Auletti for technical assistance with animal training.

