## Supplementary Materials for "Rats use darting as a strategy to navigate between reward and safety during platform-mediated active avoidance under different social contexts"

**Supplementary Figures.**

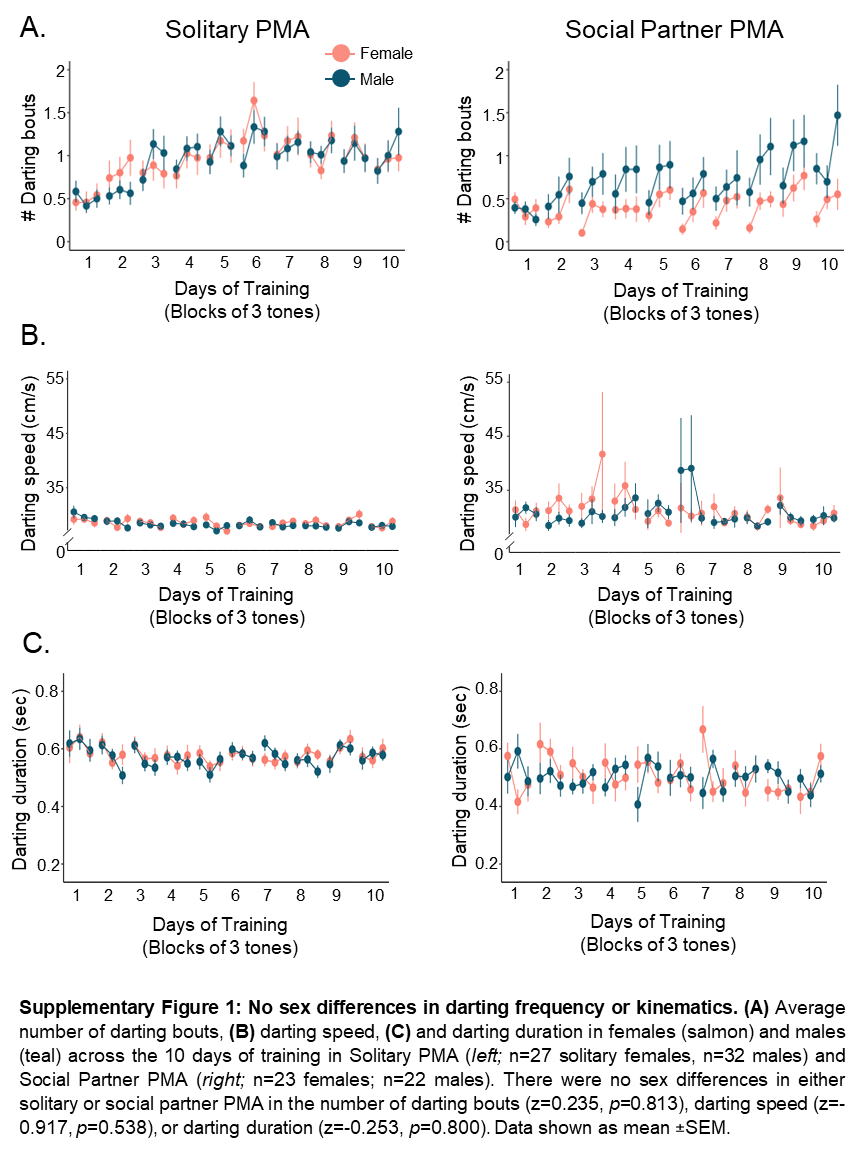

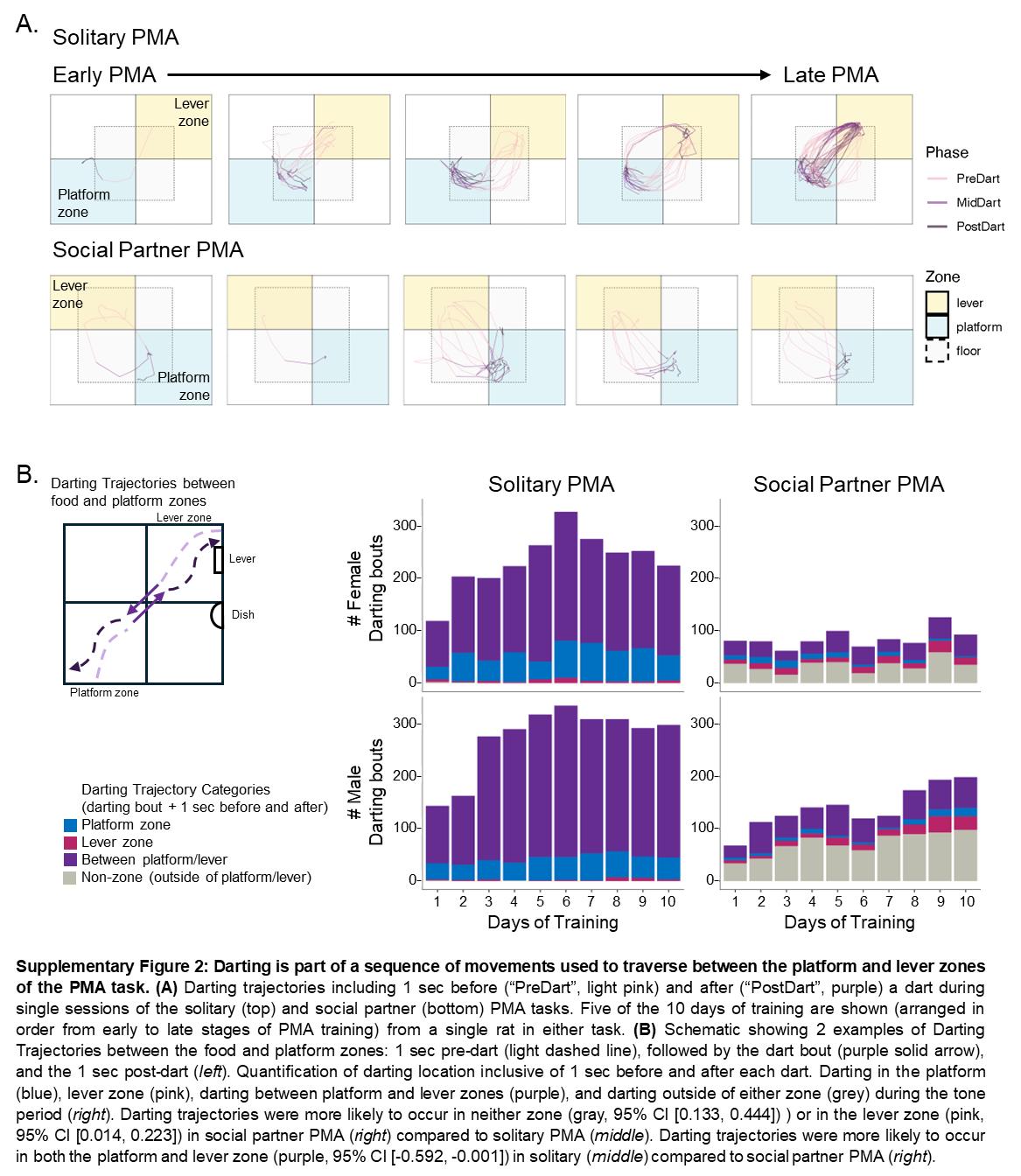

**Supplementary Tables.**

**Supplementary Table 1.**

Fixed effects parameter estimates of the multilevel negative binomial regression with Condition (social vs. solitary), Sex, Day Number, and their interactions and random effects of Subject (random intercept), Stimulus Number (random intercept) and Day Number (random slope effect by Subject), predicting the number of darting bouts per tone. All categorical variables are effect coded.

| **Parameter** | **B** | **SE** | **z** | **P** |
| --- | --- | --- | --- | --- |
| (Intercept) | -0.8841 | 0.1025 | -8.6240 | 0.0000 |
| Condition[Social] | -0.3906 | 0.0888 | -4.4004 | 0.0000 |
| Sex[Female] | 0.0209 | 0.0883 | 0.2364 | 0.8131 |
| Day Number | 0.0492 | 0.0108 | 4.5388 | 0.0000 |
| Condition[Social]:Sex[Female] | 0.0207 | 0.0883 | 0.2347 | 0.8145 |
| Condition[Social]:Day Number | -0.0038 | 0.0107 | -0.3603 | 0.7186 |
| Sex[Female]:Day Number | -0.0121 | 0.0106 | -1.1467 | 0.2515 |
| Condition[Social]:Sex[Female]:Day Number | -0.0158 | 0.0106 | -1.4960 | 0.1347 |

**Supplementary Table 2.**

Fixed effects parameter estimates of the multilevel gamma regression with Condition (social vs. solitary), Sex, Day Number, and their interactions and a random intercept effect of Subject predicting the speed of darts in cm/s. All categorical variables are effect coded.

| **Parameter** | **B** | **SE** | **z** | **P** |
| --- | --- | --- | --- | --- |
| (Intercept) | 3.404 | 0.010 | 351.753 | <0.001 |
| Condition[Social] | 0.052 | 0.009 | 5.541 | <0.001 |
| Sex[Female] | 0.004 | 0.009 | 0.479 | 0.632 |
| Day Number | -0.004 | 0.001 | -2.643 | 0.008 |
| Condition[Social]:Sex[Female] | 0.005 | 0.009 | 0.577 | 0.564 |
| Condition[Social]:Day Number | -0.002 | 0.001 | -1.313 | 0.189 |
| Sex[Female]:Day Number | <0.001 | 0.001 | -0.033 | 0.974 |
| Condition[Social]:Sex[Female]:Day Number | -0.001 | 0.001 | -0.829 | 0.407 |

**Supplementary Table 3.**

Fixed effects parameter estimates of the multilevel gamma regression with Condition (social vs. solitary), Sex, Day Number, and their interactions and a random intercept effect of Subject predicting the duration of darts in seconds. All categorical variables are effect coded.

| **Parameter** | **B** | **SE** | **z** | **P** |
| --- | --- | --- | --- | --- |
| (Intercept) | -0.6115 | 0.0127 | -48.1127 | 0.0000 |
| Condition[Social] | -0.0520 | 0.0127 | -4.0842 | 0.0000 |
| Sex[Female] | 0.0100 | 0.0127 | 0.7866 | 0.4315 |
| Day Number | -0.0011 | 0.0018 | -0.6090 | 0.5425 |
| Condition[Social]:Sex[Female] | -0.0032 | 0.0127 | -0.2533 | 0.8000 |
| Condition[Social]:Day Number | -0.0020 | 0.0018 | -1.1496 | 0.2503 |
| Sex[Female]:Day Number | -0.0028 | 0.0018 | -1.5990 | 0.1098 |
| Condition[Social]:Sex[Female]:Day Number | -0.0014 | 0.0018 | -0.7978 | 0.4250 |

**Supplementary Table 4.**

Fixed effects parameter estimates of the multilevel gamma regression with Condition (social vs. solitary), Sex, Day Number, and their interactions and a random intercept effects of Subject predicting the distance of darts in cm. All categorical variables are effect coded.

| **Parameter** | ***B*** | ***SE*** | ***z*** | ***p*** |
| --- | --- | --- | --- | --- |
| (Intercept) | 2.0820 | 0.0276 | 75.3829 | 0.0000 |
| Condition[Social] | 0.0424 | 0.0276 | 1.5353 | 0.1247 |
| Sex[Female] | -0.0028 | 0.0275 | -0.1022 | 0.9186 |
| Day Number | -0.0110 | 0.0037 | -2.9301 | 0.0034 |
| Condition[Social]:Sex[Female] | -0.0118 | 0.0275 | -0.4270 | 0.6693 |
| Condition[Social]:Day Number | -0.0033 | 0.0037 | -0.8896 | 0.3737 |
| Sex[Female]:Day Number | -0.0007 | 0.0038 | -0.1811 | 0.8563 |
| Condition[Social]:Sex[Female]:Day Number | -0.0027 | 0.0038 | -0.7204 | 0.4713 |

**Supplementary Table 5.**

Fixed effects parameter estimates of the Bayesian multilevel multinomial regression with Condition (social vs. solitary), Sex, Day Number, and their interactions and random effects of Subject (random intercept), Stimulus Number (random slope effect by Subject),) and Day Number (random slope effect by Subject), predicting the likelihood of darting bouts occurring in a given Region of Interest (Lever, Platform, or Both) compared to neither of these regions. This table includes only data from within each darting bout. All categorical variables are effect coded.

| **Comparison** | **Parameter** | **B** | **Est_Error** | **CI_2.5** | **CI_97.5** | **BF** |
| --- | --- | --- | --- | --- | --- | --- |
| Lever vs. None | Intercept | -0.7150 | 0.2933 | -1.3135 | -0.1641 | NA |
| Platform vs. None | Intercept | 1.3406 | 0.2452 | 0.8575 | 1.8269 | NA |
| Both vs. None | Intercept | -0.0631 | 0.2460 | -0.5550 | 0.4162 | NA |
| Lever vs. None | Condition[Social] | -0.7429 | 0.2762 | -1.3070 | -0.2130 | 332.3333 |
| Lever vs. None | Sex[Female] | -0.0805 | 0.2716 | -0.6093 | 0.4571 | 1.6322 |
| Lever vs. None | Day Number | -0.0308 | 0.0428 | -0.1167 | 0.0510 | 3.1863 |
| Lever vs. None | Condition[Social]:Sex[Female] | 0.3479 | 0.2693 | -0.1723 | 0.8819 | 9.3717 |
| Lever vs. None | Condition[Social]:Day Number | -0.1023 | 0.0373 | -0.1752 | -0.0287 | 288.1566 |
| Lever vs. None | Sex[Female]:Day Number | -0.0316 | 0.0367 | -0.1036 | 0.0408 | 4.1480 |
| Lever vs. None | Condition[Social]:Sex[Female]: Day Number | -0.0317 | 0.0364 | -0.1038 | 0.0389 | 4.1948 |
| Platform vs. None | Condition[Social] | -1.6700 | 0.2529 | -2.1892 | -1.1934 | Inf |
| Platform vs. None | Sex[Female] | -0.2940 | 0.2404 | -0.7628 | 0.1872 | 7.9153 |
| Platform vs. None | Day Number | 0.0140 | 0.0234 | -0.0313 | 0.0598 | 2.6469 |
| Platform vs. None | Condition[Social]:Sex[Female] | 0.1878 | 0.2387 | -0.2799 | 0.6543 | 3.7004 |
| Platform vs. None | Condition[Social]:Day Number | -0.0980 | 0.0259 | -0.1504 | -0.0487 | Inf |
| Platform vs. None | Sex[Female]:Day Number | 0.0312 | 0.0229 | -0.0138 | 0.0767 | 10.6675 |
| Platform vs. None | Condition[Social]:Sex[Female]: Day Number | 0.0064 | 0.0233 | -0.0393 | 0.0521 | 1.5510 |
| Both vs. None | Condition[Social] | -1.2371 | 0.2434 | -1.7209 | -0.7720 | Inf |
| Both vs. None | Sex[Female] | -0.3632 | 0.2382 | -0.8287 | 0.1130 | 14.9787 |
| Both vs. None | Day Number | 0.0242 | 0.0265 | -0.0271 | 0.0761 | 4.4932 |
| Both vs. None | Condition[Social]:Sex[Female] | 0.2455 | 0.2394 | -0.2261 | 0.7138 | 5.5023 |
| Both vs. None | Condition[Social]:Day Number | -0.1037 | 0.0269 | -0.1563 | -0.0508 | 3999.0000 |
| Both vs. None | Sex[Female]:Day Number | 0.0337 | 0.0255 | -0.0162 | 0.0834 | 9.6857 |
| Both vs. None | Condition[Social]:Sex[Female]: Day Number | 0.0181 | 0.0256 | -0.0321 | 0.0692 | 3.1958 |

**Supplementary Table 6.**

Fixed effects parameter estimates of the Bayesian multilevel multinomial regression with Condition (social vs. solitary), Sex, Day Number, and their interactions and random effects of Subject (random intercept), Stimulus Number (random intercept) and Day Number (random slope effect by Subject), predicting the likelihood of darting bouts occurring in a given Region of Interest (Lever, Platform, or Both) compared to neither of these regions. This table includes data from 1 second preceding and following each darting bout. All categorical variables are effect coded.

| **Comparison** | **Parameter** | **B** | **Est_Error** | **CI_2.5** | **CI_97.5** | **BF** |
| --- | --- | --- | --- | --- | --- | --- |
| Lever vs. None | Intercept | -1.7243 | 0.8031 | -3.3513 | -0.2113 | NA |
| Platform vs. None | Intercept | 1.3711 | 0.6384 | 0.1506 | 2.6472 | NA |
| Both vs. None | Intercept | 2.9288 | 0.6355 | 1.7342 | 4.2219 | NA |
| Lever vs. None | Condition[Social] | -0.2088 | 0.7026 | -1.5971 | 1.1418 | 1.5870 |
| Lever vs. None | Sex[Female] | 0.1665 | 0.6978 | -1.1967 | 1.5403 | 1.4840 |
| Lever vs. None | Day Number | 0.5230 | 0.1986 | 0.1751 | 0.9577 | 1199.0000 |
| Lever vs. None | Condition[Social]:Sex[Female] | 0.4974 | 0.6924 | -0.8493 | 1.8626 | 3.2988 |
| Lever vs. None | Condition[Social]:Day Number | -0.4509 | 0.1895 | -0.8788 | -0.1258 | 726.2727 |
| Lever vs. None | Sex[Female]:Day Number | 0.1244 | 0.1904 | -0.2187 | 0.5316 | 2.8247 |
| Lever vs. None | Condition[Social]:Sex[Female]: Day Number | -0.1987 | 0.1904 | -0.6068 | 0.1445 | 6.1471 |
| Platform vs. None | Condition[Social] | -2.1139 | 0.6286 | -3.3956 | -0.9254 | 4799.0000 |
| Platform vs. None | Sex[Female] | -0.1248 | 0.6193 | -1.3619 | 1.0628 | 1.3711 |
| Platform vs. None | Day Number | 0.4911 | 0.1875 | 0.1742 | 0.9135 | 3427.5714 |
| Platform vs. None | Condition[Social]:Sex[Female] | 0.9125 | 0.6202 | -0.2784 | 2.1332 | 13.3971 |
| Platform vs. None | Condition[Social]:Day Number | -0.5153 | 0.1864 | -0.9331 | -0.1996 | 11999.0000 |
| Platform vs. None | Sex[Female]:Day Number | 0.1802 | 0.1875 | -0.1556 | 0.5845 | 5.2305 |
| Platform vs. None | Condition[Social]:Sex[Female]: Day Number | -0.2319 | 0.1878 | -0.6350 | 0.1051 | 9.5217 |
| Both vs. None | Condition[Social] | -2.2475 | 0.6227 | -3.5139 | -1.0630 | 23999.0000 |
| Both vs. None | Sex[Female] | -0.5801 | 0.6174 | -1.8061 | 0.6144 | 4.7845 |
| Both vs. None | Day Number | 0.5023 | 0.1860 | 0.1890 | 0.9199 | 5999.0000 |
| Both vs. None | Condition[Social]:Sex[Female] | 0.7445 | 0.6149 | -0.4277 | 1.9829 | 7.9219 |
| Both vs. None | Condition[Social]:Day Number | -0.5081 | 0.1864 | -0.9245 | -0.1935 | 7999.0000 |
| Both vs. None | Sex[Female]:Day Number | 0.2136 | 0.1868 | -0.1193 | 0.6151 | 7.9519 |
| Both vs. None | Condition[Social]:Sex[Female]: Day Number | -0.1948 | 0.1865 | -0.5979 | 0.1391 | 6.2508 |

**Supplementary Table 7.**

Fixed effects parameter estimates of the multilevel negative binomial regression with Proportion of Time Spent Freezing, Condition (social vs. solitary), Sex, Day Number, and their interactions and random effects of Subject (random intercept), Stimulus Number (random intercept), Day Number (random slope effect by Subject), and Proportion of Time Spent Freezing (random slope effect by Subject) predicting the number of darting bouts per tone. All categorical variables are effect coded.

| **Parameter** | **B** | **SE** | **z** | **P** |
| --- | --- | --- | --- | --- |
| (Intercept) | 0.0106 | 0.0930 | 0.1144 | 0.9089 |
| FreezePercent | -3.7634 | 0.2700 | -13.9386 | 0.0000 |
| Condition[Social] | -0.1769 | 0.0782 | -2.2625 | 0.0237 |
| Sex[Female] | -0.0254 | 0.0779 | -0.3257 | 0.7447 |
| Day Number | 0.0012 | 0.1108 | 0.0109 | 0.9913 |
| FreezePercent:Condition[Social] | 0.2070 | 0.2627 | 0.7879 | 0.4308 |
| FreezePercent:Sex[Female] | -0.1775 | 0.2613 | -0.6791 | 0.4971 |
| Condition[Social]:Sex[Female] | -0.1065 | 0.0779 | -1.3678 | 0.1714 |
| FreezePercent:Day Number | 0.6793 | 0.3749 | 1.8120 | 0.0700 |
| Condition[Social]:Day Number | 0.0021 | 0.1091 | 0.0192 | 0.9847 |
| Sex[Female]:Day Number | 0.0562 | 0.1086 | 0.5179 | 0.6046 |
| FreezePercent:Condition[Social]:Sex[Female] | 0.2767 | 0.2616 | 1.0578 | 0.2901 |
| FreezePercent:Condition[Social]:Day Number | -0.1106 | 0.3719 | -0.2975 | 0.7661 |
| FreezePercent:Sex[Female]:Day Number | -0.8021 | 0.3722 | -2.1552 | 0.0311 |
| Condition[Social]:Sex[Female]:Day Number | 0.0683 | 0.1083 | 0.6302 | 0.5285 |
| FreezePercent:Condition[Social]:Sex[Female]:Day Number | -0.9934 | 0.3718 | -2.6718 | 0.0075 |

**Supplementary Table 8.**

Fixed effects parameter estimates of the multilevel negative binomial regression with Proportion of Time on Platform, Condition (social vs. solitary), Sex, Day Number, and random effects of Subject (random intercept), Stimulus Number (random intercept), Day Number (random slope effect by Subject), and Proportion of Time on Platform (random slope effect by Subject) predicting the number of darting bouts per tone. All categorical variables are effect coded.

| **Parameter** | **B** | **SE** | **z** | **P** |
| --- | --- | --- | --- | --- |
| (Intercept) | -0.3079 | 0.0959 | -3.2111 | 0.0013 |
| AvoidPercent | -0.8170 | 0.1144 | -7.1418 | 0.0000 |
| Condition[Social] | -0.3960 | 0.0810 | -4.8904 | 0.0000 |
| Sex[Female] | 0.0689 | 0.0807 | 0.8532 | 0.3936 |
| Day Number | 0.4077 | 0.1197 | 3.4043 | 0.0007 |
| AvoidPercent:Condition[Social] | 0.0224 | 0.1137 | 0.1967 | 0.8441 |
| AvoidPercent:Sex[Female] | -0.0869 | 0.1135 | -0.7654 | 0.4441 |
| Condition[Social]:Sex[Female] | -0.0609 | 0.0805 | -0.7565 | 0.4493 |
| AvoidPercent:Day Number | -0.0157 | 0.1621 | -0.0968 | 0.9229 |
| Condition[Social]:Day Number | 0.0153 | 0.1183 | 0.1295 | 0.8969 |
| Sex[Female]:Day Number | 0.1068 | 0.1173 | 0.9105 | 0.3626 |
| AvoidPercent:Condition[Social]:Sex[Female] | 0.0496 | 0.1132 | 0.4384 | 0.6611 |
| AvoidPercent:Condition[Social]:Day Number | -0.0732 | 0.1614 | -0.4536 | 0.6501 |
| AvoidPercent:Sex[Female]:Day Number | -0.4764 | 0.1612 | -2.9549 | 0.0031 |
| Condition[Social]:Sex[Female]:Day Number | -0.0802 | 0.1171 | -0.6852 | 0.4932 |
| AvoidPercent:Condition[Social]:Sex[Female]:Day Number | -0.1779 | 0.1612 | -1.1034 | 0.2699 |

**Supplementary Table 9.**

Fixed effects parameter estimates of the multilevel negative binomial regression with Number of Lever Presses, Condition (social vs. solitary), Sex, Day Number, and their interactions and random effects of Subject (random intercept), Stimulus Number (random intercept), Day Number (random slope effect by Subject), and Number of Lever Presses (random slope effect by Subject) predicting the number of darting bouts per tone. All categorical variables are effect coded.

| **Parameter** | **B** | **SE** | **z** | **P** |
| --- | --- | --- | --- | --- |
| (Intercept) | -1.0570 | 0.1010 | -10.4624 | 0.0000 |
| LeverCount | 0.0542 | 0.0048 | 11.3688 | 0.0000 |
| Condition[Social] | -0.2309 | 0.0886 | -2.6058 | 0.0092 |
| Sex[Female] | 0.0932 | 0.0886 | 1.0526 | 0.2925 |
| Day Number | 0.4005 | 0.1116 | 3.5885 | 0.0003 |
| LeverCount:Condition[Social] | -0.0027 | 0.0047 | -0.5718 | 0.5675 |
| LeverCount:Sex[Female] | 0.0038 | 0.0047 | 0.8061 | 0.4202 |
| Condition[Social]:Sex[Female] | -0.0311 | 0.0885 | -0.3512 | 0.7254 |
| LeverCount:Day Number | -0.0327 | 0.0057 | -5.7069 | 0.0000 |
| Condition[Social]:Day Number | -0.1854 | 0.1095 | -1.6930 | 0.0905 |
| Sex[Female]:Day Number | -0.3270 | 0.1098 | -2.9798 | 0.0029 |
| LeverCount:Condition[Social]:Sex[Female] | -0.0010 | 0.0047 | -0.2183 | 0.8272 |
| LeverCount:Condition[Social]:Day Number | 0.0072 | 0.0057 | 1.2666 | 0.2053 |
| LeverCount:Sex[Female]:Day Number | 0.0055 | 0.0057 | 0.9695 | 0.3323 |
| Condition[Social]:Sex[Female]:Day Number | -0.1513 | 0.1094 | -1.3828 | 0.1667 |
| LeverCount:Condition[Social]:Sex[Female]:Day Number | -0.0004 | 0.0057 | -0.0789 | 0.9371 |
